# Automatic segmentation of the rat brain hippocampus in MRI after traumatic brain injury

**DOI:** 10.1101/2021.08.03.454863

**Authors:** Riccardo De Feo, Elina Hämäläinen, Eppu Manninen, Riikka Immonen, Juan Miguel Valverde, Xavier Ekolle Ndode-Ekane, Olli Gröhn, Asla Pitkänen, Jussi Tohka

## Abstract

Registration-based methods are commonly used in the anatomical segmentation of magnetic resonance (MR) brain images. However, they are sensitive to the presence of deforming brain pathologies that may interfere with the alignment of the atlas image with the target image. Our goal was to develop an algorithm for automated segmentation of the normal and injured rat hippocampus. We implemented automated segmentation using a U-Net-like Convolutional Neural Network (CNN). of sham-operated experimental controls and rats with lateral-fluid-percussion induced traumatic brain injury (TBI) on MR images and trained ensembles of CNNs. Their performance was compared to three registration-based methods: single-atlas, multi-atlas based on majority voting and Similarity and Truth Estimation for Propagated Segmentations (STEPS). Then, the automatic segmentations were quantitatively evaluated using six metrics: Dice score, Hausdorff distance, precision, recall, volume similarity and compactness using cross-validation. Our CNN and multi-atlas -based segmentations provided excellent results (Dice scores > 0.90) despite the presence of brain lesions, atrophy and ventricular enlargement. In contrast, the performance of singe-atlas registration was poor (Dice scores < 0.85). Unlike registration-based methods, which performed better in segmenting the contralateral than the ipsilateral hippocampus, our CNN-based method performed equally well bilaterally. Finally, we assessed the progression of hippocampal damage after TBI by applying our automated segmentation tool. Our data show that the presence of TBI, time after TBI, and whether the location of the hippocampus was ipsilateral or contralateral to the injury explained hippocampal volume (*p* = 0.029, *p <* 0.001, and *p <* 0.001 respectively).

## 1 Introduction

In vivo Magnetic Resonance Imaging (MRI) is a key technology for tracking neuroanatomical changes in brain diseases in both experimental models and humans. Its non-invasive nature allows the design of longitudinal studies in living animals, imaging brains in three dimensions (3D) at multiple time points and using a variety of different contrasts. Region segmentation [1, 2], which systematically identifies specific regions of interest (ROIs) in 3D MRI scans, is an important component of many MRI data processing pipelines. This step is often performed manually, especially in preclinical MRI analysis. However, manual segmentation is time-consuming and suffers from inter-rater and intra-rater variability [3].

A number of automated methods exist as an alternative to manual segmentation. Typically, these methods are based on registration, where the target volume is labeled by aligning a labeled atlas to the target volume [4, 5]. The quality of the segmentation can be further improved by aligning multiple atlases to the same target volume and combining the segmentation maps. Label fusion can be accomplished by majority voting or by more complex methods, such as Similarity and Truth Estimation for Propagated Segmentations (STEPS) [6]. However, these methods depend on the quality of the registration steps. In addition, atlases typically represent healthy brains and do not account for the additional heterogeneity introduced by brain diseases or injuries such as traumatic brain injury (TBI). For example, depending on the impact force and its direction, TBI can result in large and multifocal lesions that significantly alter brain anatomy with high interindividual variability [7]. Consequently, aligning MRIs of the injured brain can be challenging, especially when the target area is in the proximity of the primary brain lesion [8].

For the segmentation of lesioned brains, Convolutional Neural Networks (CNNs) [9] provide an alternative to registration-based methods. CNNs do not directly apply manually labeled atlases to the target anatomy, but use them during training. Consequently, the information needed to segment a new sample is encoded in the neural network parameters, eliminating the need for image registration. Recently, CNNs have been successfully applied to a number of medical imaging segmentation tasks. For murine MRI, neural networks have been proposed for skull-stripping [10, 11], lesion segmentation [12, 13] and region segmentation [14].

We hypothesized that the performance of CNNs would not be affected by the presence of lesions to the same extent as registration-based methods as long as a sufficient anatomical variety of lesions is present in the training data. We segmented MRI scans acquired at different time points following experimental TBI, available from our two large preclinical animal cohorts, EpiBioS4Rx [15, 16] and EPITARGET [17, 18]. We focused on the segmentation of the hippocampus as (a) it is frequently damaged by experimental and human TBI and (b) its damage is associated with the development of posttraumatic epilepsy and cognitive impairment in both animal models and humans [19, 20]. Because hippocampal segmentation is complicated by the presence of adjacent neocortical damage and atrophy, ventricular enlargement, and hippocampal distortion, we trained MU-Net-R, a multi-task U-Net for MRI segmentation and skull stripping, and compared the results with state-of-the-art registration-based segmentation methods.

## 2 Materials and Methods

### 2.1 Materials

#### 2.1.1 EpiBioS4Rx cohort

The Epilepsy Bioinformatics Study for Antiepileptogenic Therapy (EpiBioS4Rx, https://epibios.loni.usc.edu/) is an international multicenter study funded by National Institutes of Health with the goal of developing therapies to prevent posttraumatic epileptogenesis. The 7-month MRI follow-up of the EpiBioS4Rx animal cohort has been described in detail previously [16, 15]. Here, we have analyzed the data from the UEF subcohort. We describe only the details that are important for the present study.

##### Animals

Adult male Sprague-Dawley rats (Envigo Laboratories B.V., The Netherlands) were used. They were single-housed in a controlled environment (temperature 21-23°C, humidity 50-60%, lights on 7:00 am to 7:00 pm) with free access to food and water. Severe traumatic brain injury was induced in the left hemisphere by lateral fluid percussion under 4% isoflurane anesthesia [16]. Sham-operated experimental controls underwent the same anesthesia and surgical procedures without the induction of the impact.

As summarized in Table 1, the entire cohort included 56 rats (13 sham and 43 with TBI), of which the 12 (5 sham, 7 TBI) first animals to complete follow-up were selected for manual annotation of the hippocampus. Mean impact pressure was 2.87 ± 0.82 atm in the entire cohort and 2.92 ± 1.37 atm in the manual annotation subcohort.

**Table 1:** Number of MRI scans per cohort (EpiBioS4Rx, EPITARGET) in different time points and treatment groups [traumatic brain injury (TBI), sham-operated experimental controls]. In parenthesis, the first number indicates the number of volumes used for manual annotation of the hippocampus and the second indicates the number of volumes used for manual annotation of brain masks. *Abbreviations*: d, day; TBI, traumatic brain injury.

| Timepoint | TBI | Sham |
| --- | --- | --- |
| EpiBioS4Rx |  |  |
| 2 d | 43 (7,0) | 12 (5,1) |
| 9 d | 42 (6,1) | 13 (5,1) |
| 30 d | 42 (6,1) | 13 (5,0) |
| 150 d | 40 (7,1) | 13 (5,1) |
| EPITARGET |  |  |
| 2 d | 118 (4,0) | 23 (2,2) |
| 7 d | 117 (4,2) | 23 (2,0) |
| 21 d | 117 (4,0) | 23 (2,2) |

##### MRI

Rats were imaged 2 days (d), 9 d, 1 month, and 5 months after TBI or sham surgery (Table 1) using a 7-Tesla Bruker PharmaScan MRI scanner (Bruker BioSpin MRI GmbH, Ettlingen, Germany). During imaging, rats were anesthetized with isoflurane. A volume coil was used as radiofrequency transmitter and a quadrature surface coil designed for the rat brain was used as receiver. Local magnetic field inhomogeneity was minimized using a threedimensional field map-based shimming protocol. All images were acquired using a three-dimensional multi-gradient echo sequence. A train of 13 echoes was acquired, where the first echo time was 2.7 ms, the echo time separation was 3.1 ms, and the last echo time was 39.9 ms. The image resolution was 0.16×0.16×0.16 mm^3^, the repetition time was 66 ms, the flip angle was 16°, the number of signal averages was 1, and the imaging time was 10 min 44 s. Images with different echo times were summed to produce a high signal-to-noise ratio image for segmentation and image registration.

#### 2.1.2 EPITARGET cohort

EPITARGET (https://epitarget.eu/) was a European Union Framework 7 -funded, large-scale, multidisciplinary research project aimed at identifying mechanisms and treatment targets for epileptogenesis after various epileptogenic brain insults. The 6-month MRI follow-up of the EPITARGET animal cohort has been described in detail previously [17, 18]. We describe only the details that are important for the present study.

##### Animals

Adult male Sprague-Dawley rats (Envigo Laboratories S.r.l., Udine, Italy) were used for the study. The housing and induction of left hemisphere TBI or sham injury were as described for the EpiBioS4Rx cohort. However, injury surgery was performed under pentobarbital-based anesthesia instead of isoflurane. The entire cohort included 144 rats, and images from the first 6 rats (2 sham, 4 TBI) were selected for manual annotation of the hippocampus. Mean impact pressure was 3.26 ± 0.08 atm in the entire cohort and 3.22 ± 0.02 atm in the manual annotation subcohort.

##### MRI

Imaging was performed as described for the EpiBioS4Rx cohort, except that (a) imaging was performed 2 d, 7 d, and 21 d after TBI or sham surgery (Table 1) and (b) all images were acquired with a two-dimensional multislice multigradient echo sequence. A train of 12 echoes was collected, where the first echo time was 4 ms, the echo time separation was 5 ms, and the last echo time was 59 ms. In-plane image resolution was 0.15×0.15 mm^2^, slice thickness was 0.5 mm, number of slices was 24, repetition time was 1.643 s, flip angle was 45°, number of signal averages was 4, and imaging time was 11 min 37 s. Images with different echo times were summed to produce a high signal-to-noise ratio image for segmentation and image registration.

### 2.2 Manual annotation

For outlining the ROIs, the 3D (EpiBioS4Rx) and multi-slice 2D (EPITARGET) 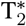-weighted MRI images were imported as NIfTI files (.nii) into Aedes 1.0 (http://aedes.uef.fi) - an in-house tool with graphical user interface for medical image analysis. Aedes is available at http://aedes.uef.fi/ and runs under MATLAB (MATLAB Release 2018b, The MathWorks, Inc.).

#### Manual segmentation of the brain mask

A trained researcher (E.H.) outlined the brain surface on 160 *µ*m-thick (EpiBioS4Rx) or 150 *µ*m-thick (EPITARGET) horizontal MRI slices, covering the entire dorsoventral extent of the cerebrum (excluding the olfactory bulbs and cerebellum). In addition, E.H. outlined the brain surface on 160 *µ*m-thick (EpiBioS4Rx) or 500 *µ*m-thick (EPITARGET) coronal brain slices to increase the accuracy of dorsal and ventral delineation of the brain surface (Supplementary Figure 1 and 2). In the EpiBioS4Rx cohort, we drew the whole brain outline for 6 scans from 6 different rats, outlining on average 33.7 ± 1.4 (range 31 – 37) horizontal slices for each MRI scan. In the EPITARGET cohort, we prepared the whole brain mask for 6 rats and the mean number of MRI slices outlined per case was 10.8 ± 0.9 (range 10 – 12).

#### Manual segmentation of the hippocampus

Outlines of the ipsilateral (left) and contralateral hippocampus were drawn by E.H. on each coronal MRI slice where the hippocampus was present (slice thickness in EpiBioS4Rx 0.16 mm and in EPITARGET 0.50 mm). In addition to the hippocampus proper and the dentate gyrus, the outlines included the fimbria fornix, but excluded the subiculum (Supplementary Figures 3 and 4). Manual annotation was performed with the help of thionin-stained coronal 30 *µ*m-thick histological sections of the same brain available at the end of follow-up, and with the Paxinos rat brain atlas [21]. In the EpiBioS4Rx cohort, we outlined the hippocampi of 15 rats (8 TBI, 7 sham) imaged at 2 d, 9 d, 30 d, and/or 5 months post-injury or sham surgery. In the EPITARGET cohort, we outlined the hippocampi of 6 rats (4 TBI, 2 sham) imaged 2 d, 7 d, and/or 21 d after TBI or sham surgery.

#### Brain mask completion

As described above, only six brain masks were manually labeled in the EpiBioS4Rx dataset and every second sagittal slice was annotated. To reconstruct complete brain masks, we first applied a binary closing operation with a hand-crafted kernel to reconstruct the brain mask, and then filled any remaining hole in the mask volume (Supplementary Figure 5). Morphological operations were implemented using the scikit-image library [22].

To generate brain masks for the complete training datasets, we trained a single 3D CNN for each dataset as described in Section 2.3, using the same overall structure and number of channels, but limiting the output to the brain mask. Using this network, we generated brain-mask labels for the remaining animals, so that our CNN could be trained on this data for both skull stripping and hippocampus segmentation.

### 2.3 CNN-based segmentation

#### 2.3.1 CNN Architecture

The architecture of our CNN (see Fig. 1) is based on MU-Net [14], which in turn was inspired by U-Net [23] and DeepNAT [24], to perform simultaneous region segmentation and skull stripping of mouse brain MRI. Our network exhibits a U-Net-like encoder/decoder structure, where the encoder and decoder branches are connected by a bottleneck layer and by skip connections between each encoder stage and the respective decoder stage. Pooling operations connect shallower blocks on the encoder branch to the deeper blocks, halving the size of the feature maps, while unpooling layers [25] connect different decoder blocks to the higher resolution ones, reversing the pooling operation.

**Figure 1:**
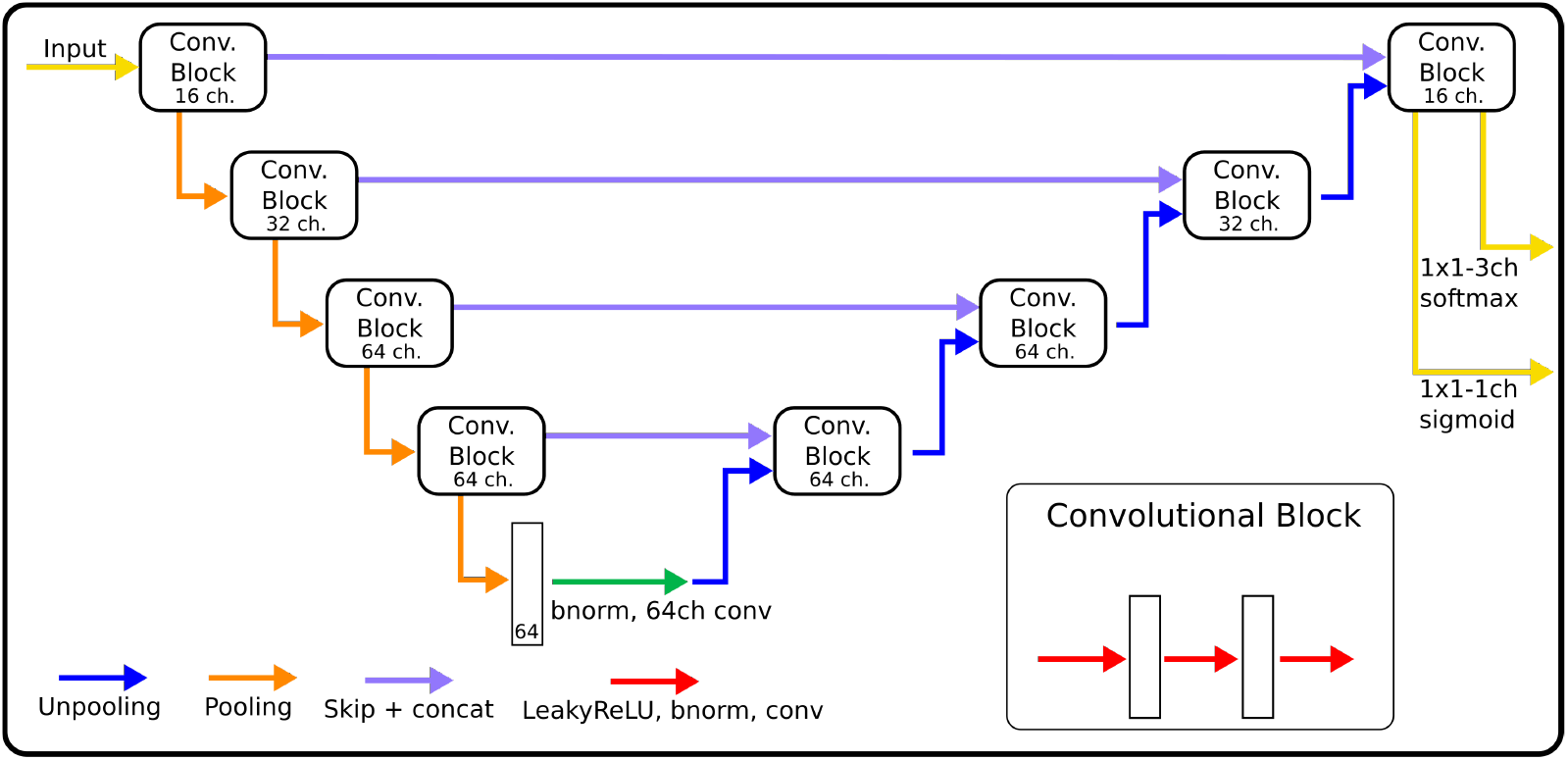
The architecture of CNNs, indicating for each block the number of channels used in each convolution. The size of convolution kernels is 3×3×3 for the EpiBioS4Rx data, and 3×3 for the EPITARGET data.

Each block consists of three iterations of Leaky ReLU activation [26], batch normalization [27] and convolution. From the shallowest to the deepest convolution block, each convolution within the block uses 16, 32, 64, and 64 channels, respectively (Fig. 1). Throughout the rest of this paper, we will refer to the network we trained for rat hippocampus segmentation as MU-Net-R.

MU-Net-R differs from MU-Net in that the size of all kernels was reduced from the 5×5 kernels in MU-Net, and that the number of kernels was reduced in the first two blocks of the neural network, whereas MU-Net used 64 kernels for each convolution.

We opted for a different choice regarding the dimension of the filters for each dataset. Since the EPITARGET 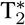 MRI data are highly anisotropic, with higher resolution on coronal slices than in the fronto-caudal direction, we chose 2D filters (3×3) in the coronal plane. This choice was based on our previous work [14] as well as [28], which indicate that a 2D approach could be preferable for the segmentation of anisotropic data. Conversely, for the network for EpiBioS4Rx 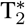 data with isotropic voxels, we preferred 3D filters (3×3×3). The total number of parameters is 428436 for the 2D EPITARGET networks and 1125716 for the 3D EpiBioS4Rx networks.

#### 2.3.2 Loss function

The loss function is an evaluation criterion minimized during the optimization of the network. Our loss function is composed of two terms, referring to the skull-stripping task (*L*_*Brain*_) and the hippocampus segmentation task (*L*_*HC*_):

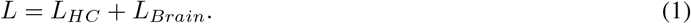

Let the prediction for label *l* at voxel *n* be defined as *p*_*ln*_, and the corresponding ground-truth value as *y*_*ln*_. For the anatomical segmentation of the hippocampus, a task including three classes (ipsilateral hippocampus, contralateral hippocampus, and background), we chose the generalized Dice loss [29] written as:

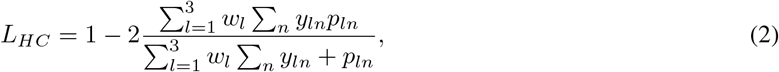

with the weight parameters *w*_*l*_ defined as *w*_*l*_ = (∑*p*_*ln*_)^*−*2^ following [29]. In practice, the weight parameters favour the hippocampus classes over the large background class. For the skull-stripping task, we used the following Dice loss term

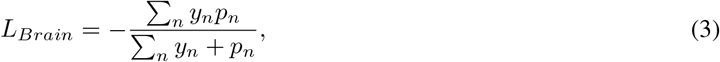

maintaining the naming conventions.

#### 2.3.3 Training

We minimized *L* with stochastic gradient descent using the RAdam optimizer [30]. RAdam is an optimizer based on Adam [31] designed to better avoid local optima, obtain more generalizable neural networks, and train in fewer epochs. We used a learning rate of 0.001, *β*_1_ = 0.9, *β*_2_ = 0.999 and no weight decay (symbols defined as in [30]). Networks and training were implemented in PyTorch and ran on a workstation with a GeForce RTX 2080 Ti GPU, 64 GB RAM and an AMD Ryzen 9 3900X 12-Core Processor. MU-Net-R networks were trained for up to 250 epochs, or until the average validation loss did not improve during the last 10 epochs.

During training, data were augmented online: each time an image was loaded, we randomly applied with a 50% probability a scaling transformation by a factor of *α*, randomly drawn from the interval [0.95, 1.05]. Each MRI volume was independently normalized to have a mean of zero and unit variance.

#### 2.3.4 Post-processing

We applied a simple post-processing procedure to each CNN-generated segmentation. We selected the largest connected component of the MU-Net-R segmentation to represent each segmented region (brain mask and hippocampus). We then filled all the holes in this component. These operations were implemented using functions from the Python module scipy.ndimage [32].

### 2.4 Registration-based segmentation

#### 2.4.1 Registration

We compared CNN-based segmentation maps with single and multi-atlas-based segmentation maps. To do this, we performed image registration using Advanced Normalization Tools [33] to facilitate the transfer of manual segmentations of the hippocampus to other brains with single- and multi-atlas approaches. Before image registration, the images were skull-stripped using FMRIB Software Library’s Brain Extraction Tool (FSL BET [34]). The masked images were then used for image registration. The brain masks for the EPITARGET dataset computed using FSL BET included marked amounts of non-brain tissue associated with the experimental traumatic brain injury, which resulted in inaccurate image registrations. To improve the brain masks, we first registered the images to one of the brain images using rigid-body and affine transformations. The FSL BET brain mask of that image was then manually refined and transformed to the rest of the brain images, resulting in more accurate brain masks and registrations.

Image registration between a template brain and a target brain volume included the computation of a rigid-body transformation, an affine transformation, and a Symmetric image Normalization (SyN) transformation. We used global correlation as the similarity metric for the rigid-body and affine transformations and neighborhood cross-correlation for the SyN transformation. The computed transforms were then applied to the template brain’s sum-over-echoes 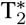-weighted image as well as its manually labeled hippocampi. All operations described in this section were performed on a 6-core AMD Ryzen 5 5600X processor.

#### 2.4.2 Single-atlas segmentation

For each template registered to a target brain, we applied the same transforms to the label map of the template, using nearest neighbor interpolation. Each measure taken for single-atlas segmentation during this work was an average between that of each individual single-atlas segmentation map for the same target brain.

#### 2.4.3 Multi-atlas segmentation

To label the hippocampus in each target volume by combining the individually-registered atlases we applied two different label fusion strategies: STEPS multi-atlas segmentation [35] and majority voting. STEPS combines multiple registered label maps by taking into account the local and global matching between the deformed templates and the target MRI volume. It does so by utilizing at the same time an expectation-maximization approach and Markov Random Fields to improve the segmentation based on the quality of the registration. We applied STEPS implementation distributed with NiftySeg [35, 6].

STEPS depends on two variables: the standard deviation of its Gaussian kernel and the number of volumes employed. Given the limited size of our dataset, and to reduce the risk of overfitting, we chose the standard deviation found in our previous work [14] as a result of a grid search between volumes aligned using diffeomorphic registration. When labeling each volume, we used all available registered atlases in label fusion for both STEPS and majority voting. The majority voting refers to choosing the most commonly occurring label among all registered atlases for each voxel.

For all atlas-based methods, i.e. single-atlas, STEPS, and majority voting, we only used as atlases those scans acquired at the same time-point.

### 2.5 Evaluation metrics

We compared the segmentation maps to the manually annotated ground truth using the following metrics: Dice score, Hausdorff 95 distance, volume similarity, compactness score, precision, and recall.

#### Dice score

The Dice score [36] is a measure of the overlap between two volumes and is defined as:

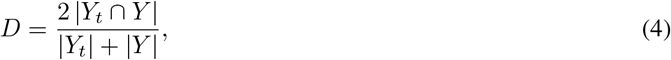

where *Y* is the automatic segmentation and *Y*_*t*_ the ground truth. A score of 1 corresponds to a perfect overlap and a score of 0 to a complete absence of overlap.

#### Hausdorff 95

We evaluated the 95th percentile of the Hausdorff distance [37], which in turn refers to the magnitude of the largest segmentation error of the prediction when compared to the ground truth. We refer to this metric as HD95 and measure it in millimeters. HD95 was calculated using MedPy [38].

#### Volume similarity

We measured the volume similarity (VS) between prediction and ground truth, following the definition provided by [39]. Unlike the Dice score, VS does not depend on the overlap between the two regions, and only depends on their volumes:

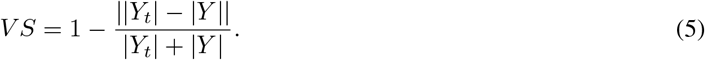

#### Compactness score

Compactness is defined as the ratio between area and volume [40]:

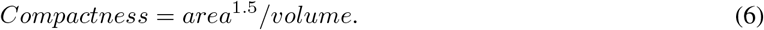

We define a compactness score to indicate how close the compactness of the generated segmented ROI, denoted by *C*, is to the compactness of the ground truth, denoted by *C*_*GT*_. The compactness score (CS) is defined:

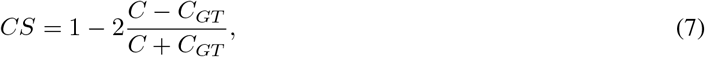

where *CS* = 1 indicates an identical compactness, and lower values indicate the two regions display a different ratio between surface and volume. To calculate the compactness, we used code from [12].

#### Precision and recall

Precision and recall evaluate respectively the ratio between true positives and the total number of positive predictions, and true positives and ground truth size. As such, increasing the number of false positives reduces the precision, and increasing the number of false negatives reduces recall. Both metrics vary between 0 and 1, and are defined as:

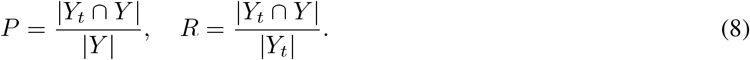

### 2.6 Cross-validation

We used cross-validation to evaluate CNN-based segmentation maps as well as the registration-based segmentation maps. For both the EpiBioS4Rx and the EPITARGET datasets, we applied 6-fold cross-validation. The labeled samples from the EpiBioS4Rx dataset were divided into 6 folds, where each fold contained two animals. Likewise, for the EPITARGET dataset, we defined 6 folds, each containing one animal.

For registration-based methods, the brain of each animal was registered to the brains of those animals that did not belong to the same fold. Registrations were performed within each time point. Thus, the brain of each of the 12 animals was registered to 10 other brains at each of the 4 time points, which would have resulted in 480 image registrations. However, images for one brain at two time points were missing, reducing the total to 440 image registrations. For the EPITARGET dataset, the brain of each of the 6 animals was registered to the 5 other brains at each of the 3 time points, resulting in 90 image registrations.

To train MU-Net-R the training set was further randomly divided into 6 folds, using one fold as validation data during the training loop for early stopping as described above. In this way, we trained an ensemble of six networks, one for each validation fold. The final prediction for the test fold was the majority voting prediction from all networks generated from the same training set. The strategy for the EPITARGET data was identical, with the only difference that in this case each fold was populated by only one animal.

We evaluated the differences in average values of evaluation metrics between different segmentation methods via a studentized paired permutation test using t-statistic as the test statistic [41]. We used 10,000 iterations for the permutation tests and applied Bonferroni’s correction for multiple comparisons.

### 2.7 Visual evaluation

After segmenting all hippocampi from the EpiBioS4Rx dataset (56 animals across 4 timepoints) with MU-Net, our annotator (E.H.) evaluated every segmented volume according to the following procedure: For each volume, one coronal slice at a time was selected, starting caudally and proceeding in the rostral direction. For each slice, the annotator was asked to input four numbers, corresponding to an evaluation for the dorsal and ventral parts of the left and right hippocampus. The evaluation scale, inspired by [42], was as follows:

1. Acceptable ’as is’
2. Minor differences. Minor edits necessary. A small number of voxels, or less than 20% of the area
3. Moderate edits required, 20–50% of the area would need to be changed
4. Major edits required, >50% of the area would need to be manually edited
5. Gross error, no resemblance to the anatomical structure

We simultaneously displayed the unlabeled MRI slice side-by-side with the same MRI slice overlaid with the ipsilateral hippocampus highlighted in red, and the contralateral one in blue. The volumes were presented to the annotator in a randomized order.

In addition to every labeled slice, we evaluated two additional slices in each direction, rostrally and caudally, to allow for the detection of hippocampal regions erroneously labeled as background. As the number of misclassified voxels was small in these cases we classified errors in these slices with a score of 2, to avoid introducing a bias because of the choice of performing this evaluation on coronal slices.

### 2.8 Statistical analysis of the hippocampal volumes

Using the trained CNNs, we labeled every MRI volume in our datasets (220 for EpiBioS4Rx and 424 for EPITARGET). As a demonstration of the applicability of the segmentation, we studied the effects of TBI on hippocampal volume through time and across both hippocampi using a repeated measures linear model, implemented using the linear mixed model function in IBM SPSS Statistics for Windows, version 26.0 (SPSS Inc., Chicago, IL, United States). Every variable was considered as a fixed effect and we assumed a diagonal covariance structure of the error term. Let *t* indicate the time point in days as a scalar variable, *R* be defined such so that *R* = 1 indicates the ipsilateral hippocampus and *R* = 0 the contralateral hippocampus. Additionally, let *B* = 1 indicate the presence of TBI, with *B* = 0 indicating sham animals, and let *E* be the error term. Then, our linear model for the volume *V* can be written as:

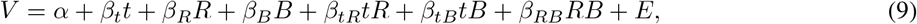

where *α, β*_*i*_ are parameters of the model.

## 3 Results

We automatically annotated every manually-labeled image in the EpiBioS4Rx and EPITARGET datasets using multiatlas segmentation (STEPS and majority voting), single atlas segmentation, and MU-Net. On a qualitative level, both multi-atlas methods and MU-Net-R showed visually convincing segmentation maps, while single-atlas segmentation resulted in the lowest-quality results (Figures 2 and 3). Where the hippocampus was markedly displaced by the injury, we noticed that registration-based methods could mislabel the lesioned area as hippocampus, as displayed in Figure 3.

**Figure 2:**
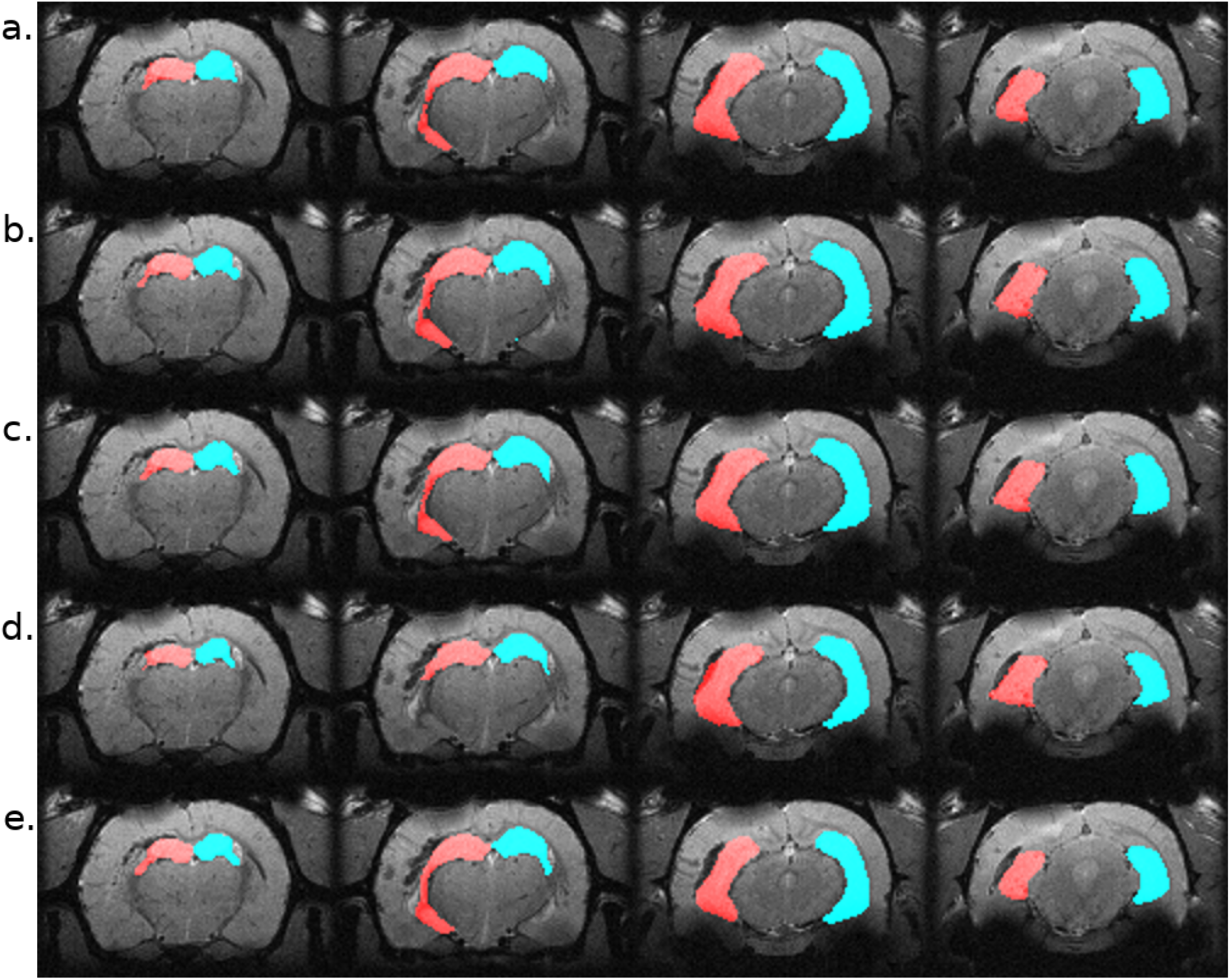
Segmentation maps in 4 representative slices from a randomly selected animal from the EpiBioS4Rx dataset, at the 2-days timepoint. Segmentation maps were obtained with: a. MU-Net; b. STEPS, c. Majority voting, d. Single-atlas segmentation. e. Displays the ground-truth segmentation. From left to right, slices are located at approximately *−*2.2, *−*3.3, *−*5.0, *−*6.2 mm from bregma. Red: hippocampus ipsilateral to the lesion; blue: hippocampus contralateral to the lesion.

**Figure 3:**
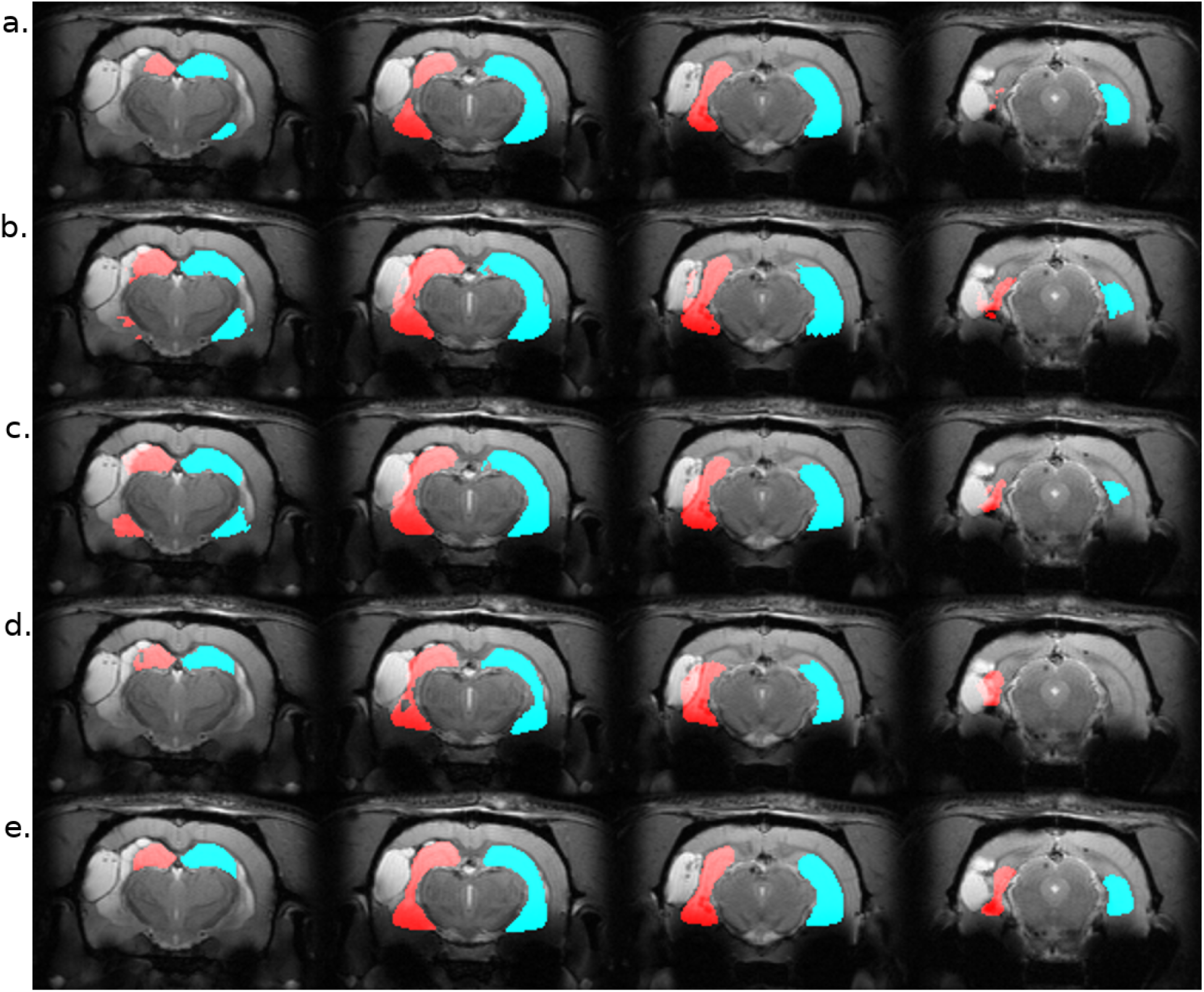
Segmentation maps in 4 representative slices from a randomly selected animal from the EPITARGET dataset, at the 21-days timepoint. Maps are obtained with: a. MU-Net; b. STEPS, c. Majority voting. d. Single-atlas segmentation. e. Displays the ground-truth segmentation. From left to right, slices are located at approximately *−*3.5, *−*4.5, *−*5.5, *−*6.5 mm from bregma. Red: hippocampus ipsilateral to the lesion; blue: hippocampus contralateral to the lesion. Note how in this case all methods except MU-Net-R mislabel a portion of the lesion as hippocampus.

### 3.1 EpiBioS4Rx segmentation

MU-Net obtained excellent segmentation evaluation scores in both hemispheres as illustrated in Fig. 4. For the ipsilateral and contralateral hippocampus MU-Net achieved, respectively, average Dice scores of 0.921 and 0.928, HD95 distances of 0.30 mm and 0.26 mm, precision of 0.935 and 0.936, recall of 0.909 and 0.921, VS of 0.968 and 0.971, and CS of 0.974 and 0.979.

**Figure 4:**
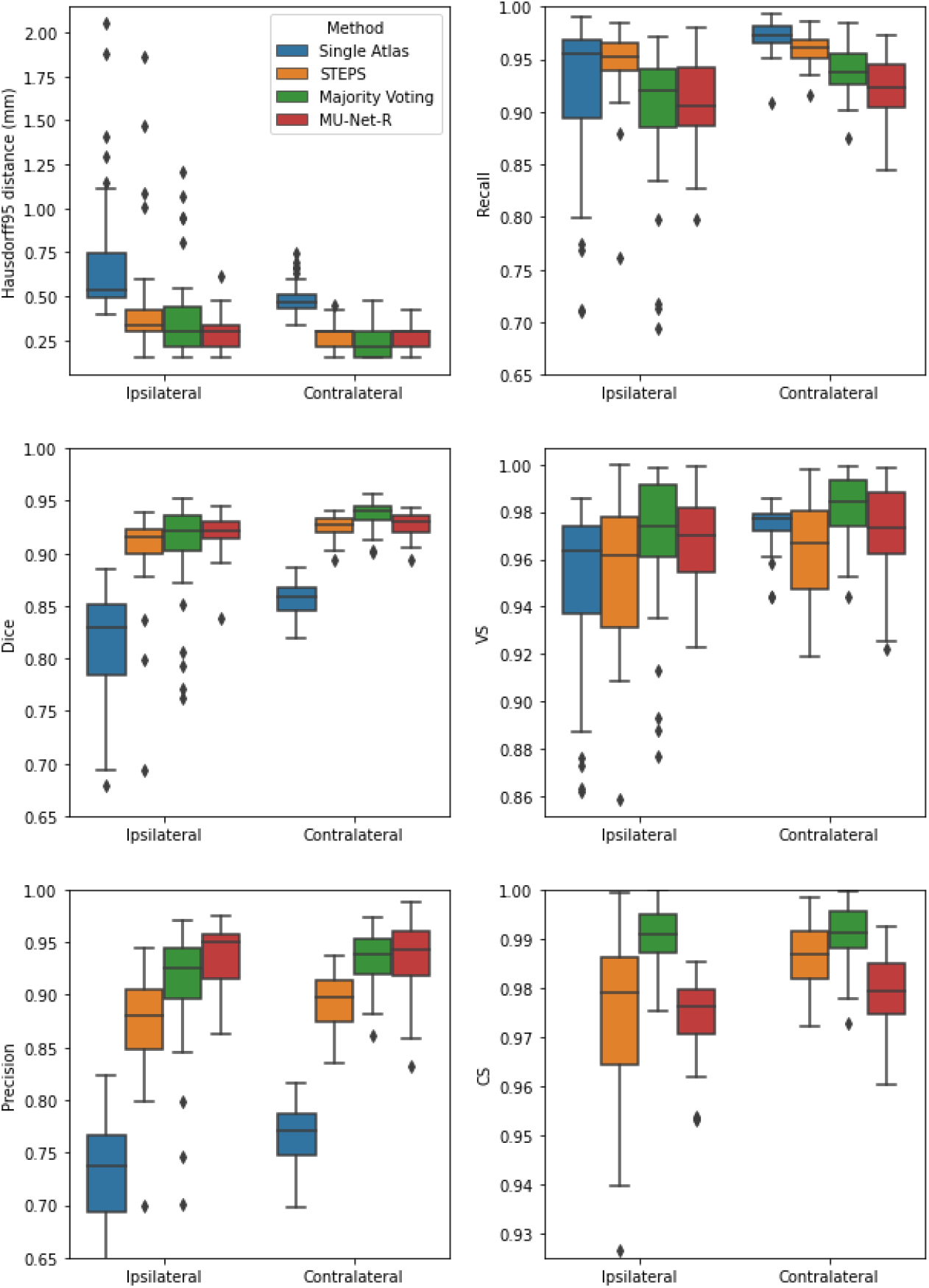
Box plots of all measured quality metrics for the contralateral and ipsilateral hippocampus in the EpiBioS4Rx dataset. Single atlas CS measures (not displayed) average at 0.60 for both hippocampi with a standard deviation of 0.01. Contrary to single-atlas segmentation, MU-Net-R and multi-atlas methods achieve human-level performance.

Quantitatively, we observed a stark difference in performance between single-atlas segmentation and all other methods in terms of Dice score, Precision, HD95, and CS (*p <* 0.001), with single-atlas segmentation performing as the least effective method. The performance measures of all other methods were excellent, with several minor differences for each metric. In terms of HD95 distance, the best performing method was MU-Net on the ipsilateral hemisphere, and majority voting in the contralateral one (*p <* 0.05). The same pattern held for the Dice score. MU-Net achieved the highest precision in the ipsilateral hippocampus (*p <* 0.05). We found no significant difference between the precision of MU-Net and that of majority voting in the contralateral one (*p >* 0.7). The precision of both methods was markedly higher than STEPS and single atlas (*p <* 0.05). As an exception to the general trend, single-atlas segmentation showed the highest value of the recall metric in the contralateral hippocampus (*p <* 0.001), while STEPS outperformed it in the ipsilateral one (*p <* 0.02). We found again no significant difference (*p >* 0.8) in the VS scores for majority voting and MU-Net in the ipsilateral hippocampus, with these two methods achieving the highest VS scores (*p <* 0.05). In contrast, majority voting achieved higher VS in the contralateral side (*p <* 0.0005). Majority voting also better preserved the compactness properties of the hippocampal shape, achieving the highest CS among all the methods (*p <* 0.0005).

### 3.2 EPITARGET segmentation

For the EPITARGET dataset, we observed similar pattern of the segmentation evaluation metrics to the ones of EpiBioS4Rx, bilaterally recording good performance metrics for MU-Net-R (see Fig. 5). We measured, respectively, for the ipsilateral and contralateral hippocampus, Dice scores of 0.836 and 0.838, HD95 distances of 0.46 mm and 0.43 mm, precision of 0.897 and 0.881, recall of 0.787 and 0.804, VS of 0.928 and 0.946, and CS of 0.992 and 0.991.

**Figure 5:**
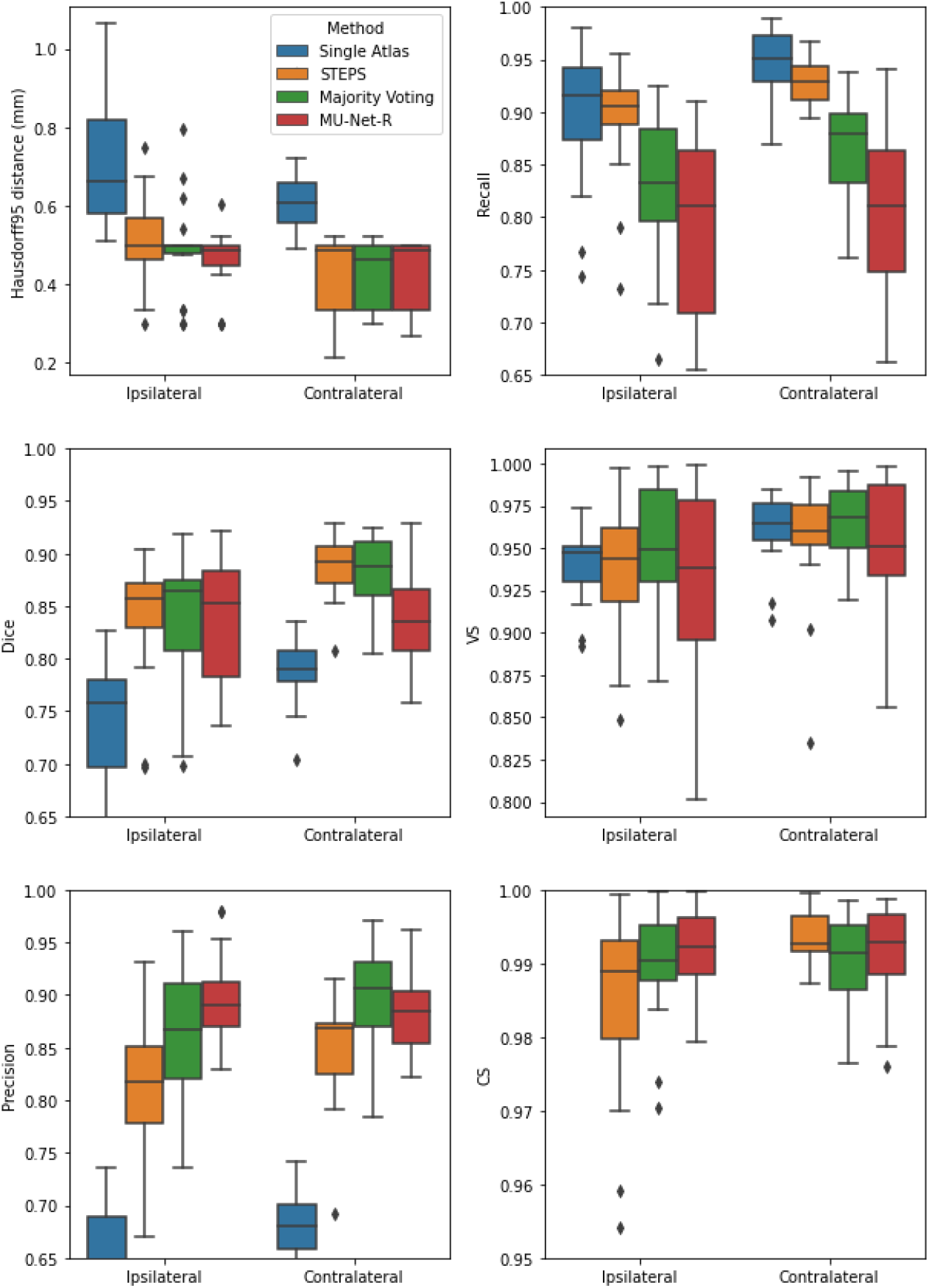
Box plots of all measured quality metrics for the contralateral and ipsilateral hippocampus in the EPITARGET dataset. Single atlas CS measures (not displayed) average at 0.67 for both hippocampi with a standard deviation of 0.02. While MU-Net-R and multi-atlas methods still outperform single-atlas segmentation, in this anisotropic dataset, training MU-Net-R with a smaller dataset, we register a lower performance compared to the EpiBioS4Rx results.

In terms of HD95 distance, on the contralateral side, we only saw a significant difference between the under-performing single-atlas method and all other methods (*p <* 0.0005). For the ipsilateral side, we additionally observed a small advantage for MU-Net-R over STEPS (*p <* 0.05). Similarly, there was no significant Dice score difference in the ipsilateral hippocampus between all high performing methods. In the contralateral one, both STEPS and majority voting achieved higher Dice scores than in the lesioned hemisphere (*p <* 0.001), achieving a higher Dice score than MU-Net-R (*p <* 0.03), which in contrast did not perform in a statistically different way (*p >* 0.7) between the two hemispheres. We measured higher precision for majority voting and MU-Net-R compared to all other metrics (*p <* 0.001) and no significant difference between the two in both the contralateral and ipsilateral hippocampus (*p >* 0.1). We observed higher recall bilaterally for single atlas segmentation compared to all other methods (*p <* 0.01), with the exception of STEPS on the ipsilateral hemisphere, where the difference was not significant (*p >* 0.2). No significant difference was also detected for VS between the different methods (*p >* 0.1) and for CS on the contralateral hippocampus. Conversely, MU-Net-R performed better than STEPS on the ipsilateral hippocampus (*p <* 0.05).

### 3.3 Inter-hemispheric differences

All segmentation methods obtained better results on the contralateral hippocampus. However, the inter-hemispheric differences for each metric were the smallest for MU-Net-R (Fig. 6), calculated as the difference in each metric for each brain between the segmentation of the ipsilateral and the contralateral hippocampus. MU-Net-R achieved significantly smaller inter-hemispheric differences in all metrics than other methods (maximal *p <* 0.02) with the exception of recall, where both MU-Net-R and STEPS perform better than all other methods (*p <* 0.03), and CS, where majority voting compares favorably to both STEPS and MU-Net-R (maximal *p <* 0.02).

**Figure 6:**
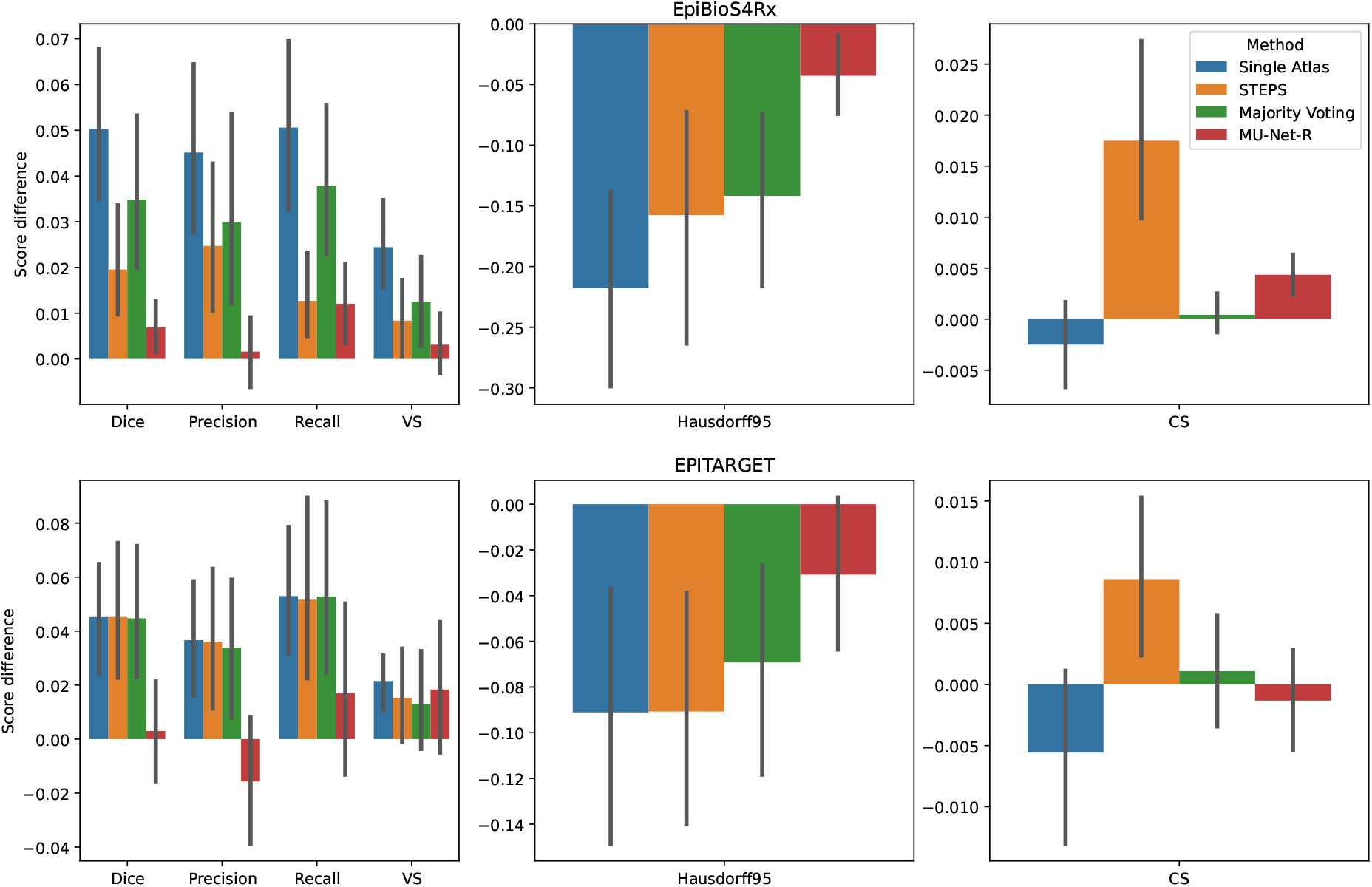
Mean inter-hemispheric differences for all evaluation metrics in single-atlas, STEPS, Majority Voting and MU-Net-R segmentation. Error bars correspond to 95% bootstrapped confidence intervals for the mean. MU-Net-R minimizes inter-hemispheric performance differences across all metrics and on both the EpiBioS4Rx and EPITARGET datasets, with the exception of CS on the EpiBioS4Rx dataset.

For the EPITARGET dataset and for all evaluation metrics, we observed a smaller average amplitude of the inter-hemispheric differences for MU-Net-R than for other segmentation methods (Fig. 6), with a single exception of CS for majority voting. However, differences were statistically significant only for the Dice score (maximal *p <* 0.02), where MU-Net-R demonstrated a stable performance between the two hemispheres. In this case, the average Dice score difference of MU-Net-R was 0.003 with a standard deviation of 0.040, while all other methods displayed an average difference of 0.045 and standard deviations of at least 0.044.

### 3.4 Segmentation time

The inference time of MU-Net-R was lower than one second per volume. The training of one ensemble of MU-Net-Rs with early stopping required on average 124 minutes for EpiBioS4Rx and 64 minutes for EPITARGET. Registering a single volume pair required approximately 40 minutes for EpiBioS4Rx volumes and 6 minutes for EPITARGET volumes. After all volumes were registered, applying majority voting and STEPS label fusion required approximately 10 seconds per target volume. Thus, multi-atlas segmentation with 10 atlases required 400 minutes for EpiBioS4Rx and multi-atlas segmentation with 5 atlases required 30 minutes for EPITARGET.

### 3.5 Visual evaluation

We visually evaluated 33136 slices from the 220 volumes in the EpiBioS4Rx dataset as described in section 2.7. The quality of the MU-Net-R segmentation in the ventral and dorsal aspects of both hippocampi was evaluated on a scale of 1 to 5, with 1 representing an accurate segmentation and 5 indicating a complete lack of resemblance to the anatomical structure, as outlined in Section 2.7. In the vast majority of cases, the reported score was 1, with a small reduction in accuracy for the ipsilateral hippocampus (Fig. 7). Overall, we found that 88.00% hippocampal regions were labeled as 1, 10.64% labeled as 2, 1.09% labeled as 3, 0.22% labeled as 4, and 0.05% labeled as 5. As illustrated in Fig. 8, the accuracy of the segmentation was the lowest in the most rostral and most caudal coronal slices. Supplementary Figure 6 provides examples of segmented slices of each score.

**Figure 7:**
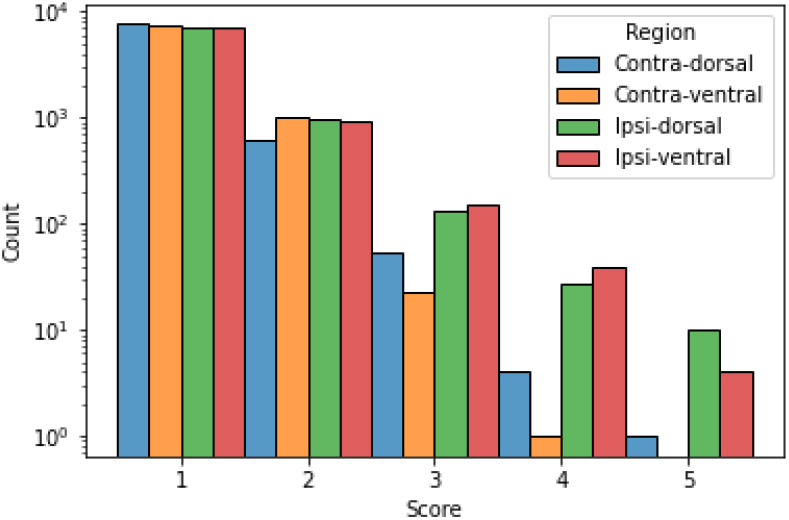
Qualitative score distribution for both hippocampi, divided for the dorsal and ventral aspects of the contralateral and ipsilateral hippocampus. Counts are reported on a logarithmic scale. The overwhelming majority of hippocampal regions were labeled as requiring no corrections (1), followed by regions requiring minor corrections only (2).

**Figure 8:**
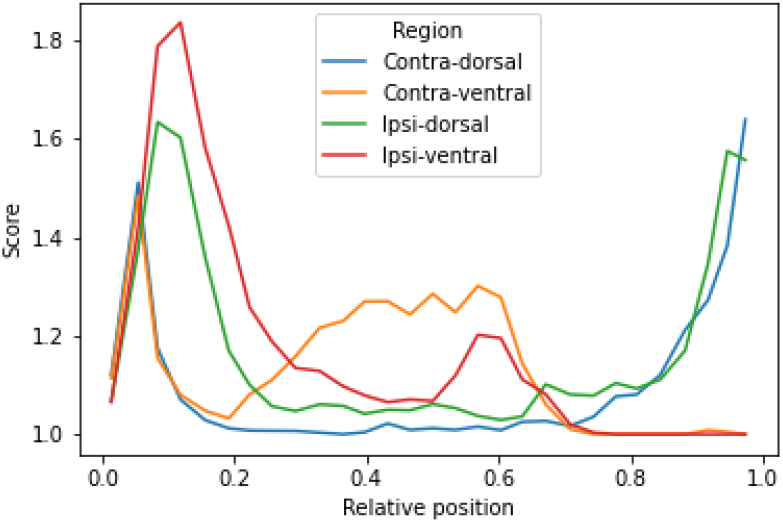
Average qualitative scores as a function of the relative position of the slice across the hippocampus, with 0 indicating the most caudal and 1 the most rostral coronal slice. Averages were obtained by dividing the interval in 30 bins. The vast majority of inaccuracies are located in the most rostral and caudal slices.

### 3.6 Hippocampal volumes

Using MU-Net-R, we annotated every scan in both the EpiBioS4Rx and EPITARGET dataset and modeled hippocampus volumes as outlined in Section 2.8. As displayed in Fig. 9, when comparing sham and TBI rats in EpiBioS4Rx we found that all included factors were statistically significant in explaining the volume: lesion status (sham or TBI), timepoint, and ROI, as well as their pairwise interaction terms. With the exception of the presence of lesions (*p* = 0.029), all other p-values were smaller or equal to 0.001, for both single factors and interaction terms. The same was true for the EPITARGET dataset, where all factors were highly significant (*p <* 0.002).

**Figure 9:**
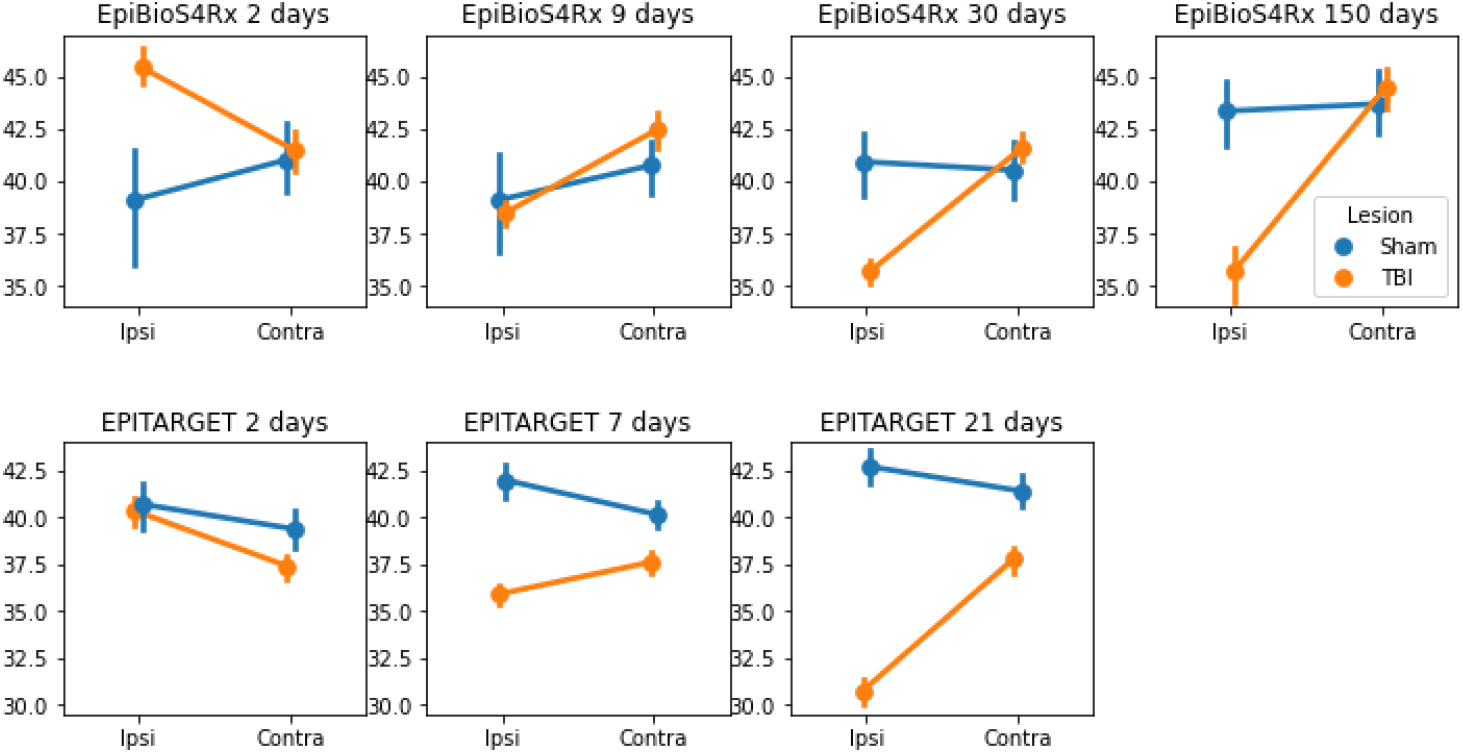
Comparison between the hippocampal volumes (mm^3^) for each dataset and time point between sham and TBI animals. Note the different dynamics in ipsilateral hippocampal volume changes between the EpiBioS4Rx and EPITARGET animals, particularly at 2 days post-injury. The rats in the EpiBioS4Rx cohort were anesthetized with 4% isoflurane at the time of injury. The rats in the EpiTARGET cohort were anesthetized with pentobarbital-based anesthesia cocktail, which apparently reduced the acute post-impact seizure-related swelling better than isoflurane.

Supplementary Table 1 provides the complete SPSS output, including *β* coefficients, p-values, *F* and *t* statistics.

## 4 Discussion

We designed and trained a CNN-based approach (MU-Net-R) to hippocampal segmentation in rat brain MRI after TBI. We quantitatively evaluated MU-Net-R-based segmentation and compared it with single- and multi-atlas registration-based segmentation methods through a variety of metrics (Dice score, volume similarity, Hausdorff distance, compactness score, precision, and recall). These evaluations demonstrated that MU-Net-R achieved state-of-the-art performance in terms of segmentation quality while reducing the bias for healthy anatomy, typical of registration-based methods, and with a marked reduction in the segmentation time compared to registration-based methods.

Single-atlas segmentation displayed the least satisfactory results, with a marked decrease in segmentation quality in terms of precision, Dice score, and HD95 distance compared to all other methods. Conversely, STEPS, majority voting, and MU-Net-R displayed excellent performance, with Dice coefficients compatible with human inter-rater agreement [3, 43, 44]. Differences in each performance metric between these methods were overall small, and different metrics would indicate a different preference. Thus, a global preference between MU-Net-R, STEPS, and majority voting did not emerge simply from these measures.

In contrast, as illustrated in Fig. 6 and 6, compared to the other segmentation methods, MU-Net-R segmentation resulted in a marked reduction in the differences for each metric between the hippocampi ipsilateral and contralateral to the lesion. A small bias was still measured, both quantitatively and qualitatively. We detected a larger number of slices evaluated with scores of 3 or higher for the ipsilateral hippocampus. This difference was likely owed to the small size of the training set; the presence of lesions manifesting in different shapes and sizes implies a larger degree of variability that requires more data to be captured. Taking this small difference into account, the vast majority of slices required no correction regardless of ROI, with only 2% of slices requiring moderate or larger edits.

A reduced performance of registration-based segmentation methods in the presence of lesions has been documented in human MRI [8]. Here, as an additional contribution of our work, we quantified and documented this difference in this example of rat brain MRI with TBI according to six different metrics. This reduced performance might imply the presence of random or systematic errors in terms of the shape or positioning of the segmentation of the ipsilateral hippocampus.

In analogy with our previous work [14], MU-Net-R was characterized by comparatively high precision and lower recall. While this difference was not large (compared to the drop in precision for single-atlas segmentation) it may still indicate a bias for the background class, which may not be entirely corrected by the weighting parameters introduced by the generalized Dice loss. Interestingly, the large drop in precision for single-atlas segmentation was largely corrected by combining multiple atlases through majority voting at the price of a lower recall, although this precision remained lower than that measured for MU-Net-R.

The inference time for MU-Net-R was markedly faster than that of registration-based methods, requiring less than one second. As for training, using the early stopping strategy described in Section 2.3.3, with this dataset size, training one ensemble of CNNs required a comparable amount of time to that required by registration, with the added benefit of obtaining a network that can then be used to quickly label a large number of new volumes. For this reason, our method is also indicated when processing large datasets, or where time constraints could be relevant, for example, in an online system designed to quickly provide a segmentation map to the user. The training time is especially low in the case of the EPITARGET data, where the number of training volumes is smaller, and the limited resolution further reduces the amount of information present in each data sample. For the same reasons, results on this dataset were less statistically meaningful, although they reflect the same general patterns as the EpiBioS4Rx data.

Consistently with [45], we observed a very small difference between Dice scores of STEPS and majority voting multi-atlas segmentation, of 0.012 on the contralateral hippocampus and 0.003 on the ipsilateral one. Thus, STEPS and majority voting appear to be largely equal according to this metric. In contrast, in our previous work [14] we measured a stronger and statistically significant difference for the segmentation of the healthy hippocampus between the two methods. This resulted in a Dice score difference of 0.055 in favor of STEPS. Two possible reasons why this difference is not observed in our case may be found in the different MRI setup, featuring a different coil and generating visibly different images. Furthermore, it could be a consequence of having performed diffeomorphic registration using a different method, FSL FNIRT [46], while the present work implemented registration tools from ANTs [33].

Thanks to the segmentation maps generated by the CNN, we could quickly label the entire EpiBioS4Rx dataset. Then, to exemplify a data analysis pipeline, using a simple regression model we identified an effect of the presence of the lesion on the volume of both hippocampi, both in general and combined with timepoint and ROI. The presence of TBI was an important predictor of hippocampal volume both as a single factor and in combination with timepoint and ROI, indicating that the volume of both the ipsilateral and contralateral hippocampus changed as a consequence of TBI.

Even though the size of the entire dataset is larger than in most preclinical studies so far, the small training dataset sizes constitute one of the limitations of our work. It is reasonable to believe that larger training datasets would further boost the performance of our neural network. Another unsolved problem remains the high degree of specialization for the specific MRI setup. While this can be attacked with a variety of methods, e.g. transfer learning [47], this still represents an open issue for CNNs. In contrast, registration-based methods can be more flexible, by using an inter-modality cost function [4].

With our work, we demonstrated how by replacing registration-based methods with CNNs we can at the same time increase the speed of segmentation, and obtain more reliable delineations of the ROIs in the presence of anatomical alterations. While we limited our analysis to one specific architecture, we do not expect this robustness to anatomical alternations to be a feature unique to our neural network, or even restricted to the U-Net-like architectures commonly applied in medical imaging. As CNNs eliminate the reliance on registration, replacing it with the encoding of knowledge in the parameters of the network itself, and given their intrinsic properties of spatial invariance, we expect segmentation CNNs, in general, to be more robust to anatomical alterations compared to registration-based methods.

## Supporting information

Supplementary images

## 5 Declarations

### 5.1 Data availability statement

The code used in our work is freely available under the MIT license at https://github.com/Hierakonpolis/ MU-Net-R. The MRI data is stored on the UEF servers and will be made available upon request.

### 5.2 Ethics statement

All experiments were approved by the Animal Ethics Committee of the Provincial Government of Southern Finland and were performed in accordance with the guidelines of the European Community Directives 2010/63/EU.

### 5.4 Credit authorship contribution statement

**Riccardo De Feo:** Methodology, Software, Formal analysis, Investigation, Writing - original draft **Elina Hämäläinen:** Data curation, Methodology. **Eppu Manninen:** Software, Investigation. **Riikka Immonen:** Investigation. **Juan Miguel Valverde:** Methodology, Writing - review and editing. **Xavier Ekolle Ndode-Ekane:** Data curation, Formal analysis, Writing - original draft **Olli Gröhn:** Conceptualization. **Asla Pitkänen:** Conceptualization, Methodology, Writing - original draft. **Jussi Tohka:** Conceptualization, Methodology, Writing - original draft

## 6 Acknowledgments

This study was supported by the Medical Research Council and Natural Science and Engineering Research Council of the Academy of Finland (Grants 272249, 273909, and 2285733-9 to AP, 316258 to JT) and by the European Union’s Seventh Framework Programme (FP7/2007-2013) under grant agreement n°602102 (EPITARGET)(AP), the National Institute of Neurological Disorders and Stroke (NINDS) Centres without Walls [grant number U54 NS100064](AP), The Sigrid Juselius Foundation (AP), by grant S21770 from the European Social Fund, grant 65211916 from the North Savo Regional Fund, from the European Union’s Horizon 2020 Framework Programme (Marie Skłodowska Curie grant agreement #740264 (GENOMMED)) (JV), and by the Alfred Kordelin Foundation (EM).

