## Supplementary images for "Automatic segmentation of the rat brain hippocampus in MRI after traumatic brain injury"

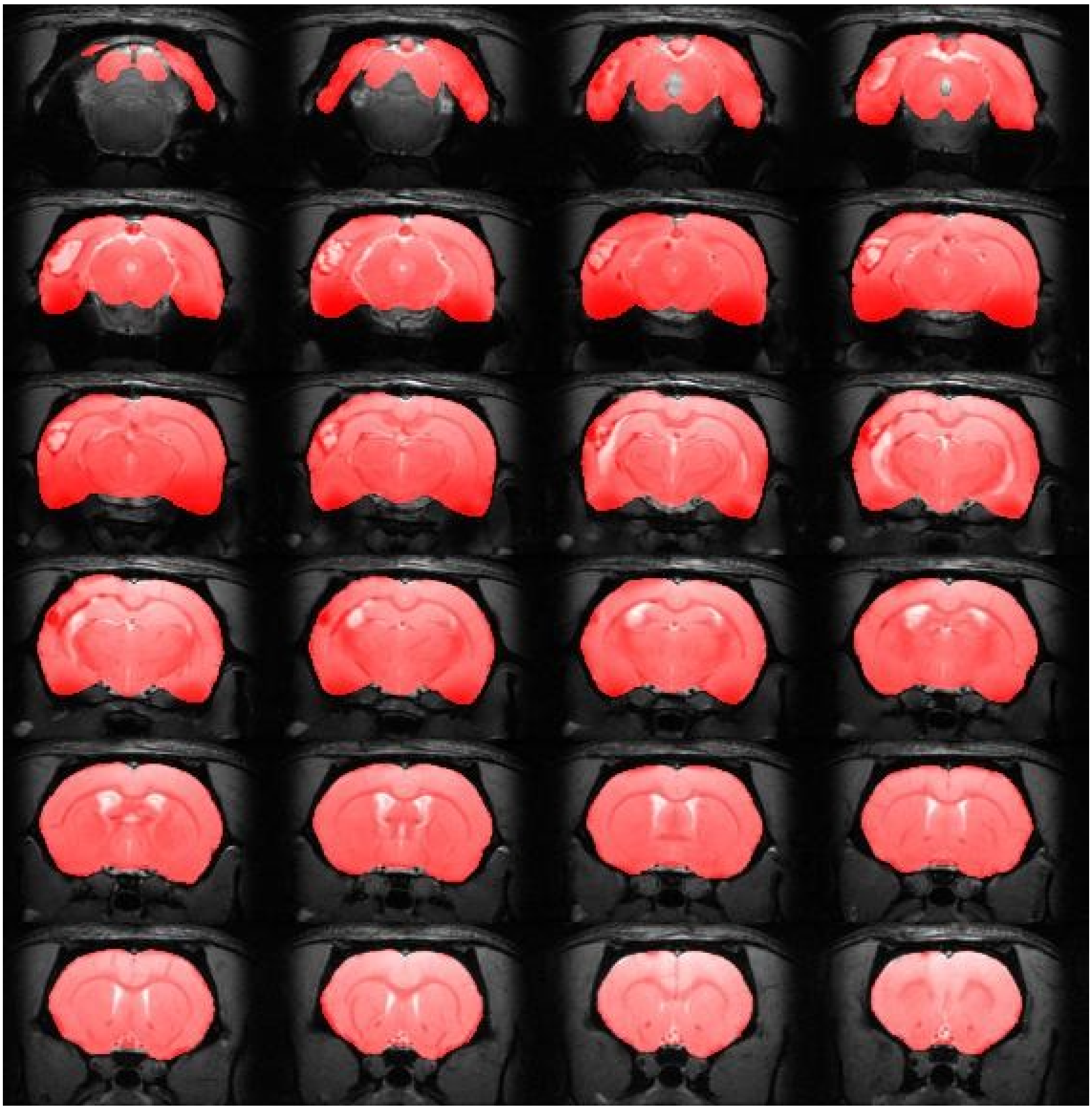

*Figure 1: Manual annotation of the brain mask in an EPITARGET rat, MRI scan acquired 21 days after TBI, displaying every annotated coronal slice.*

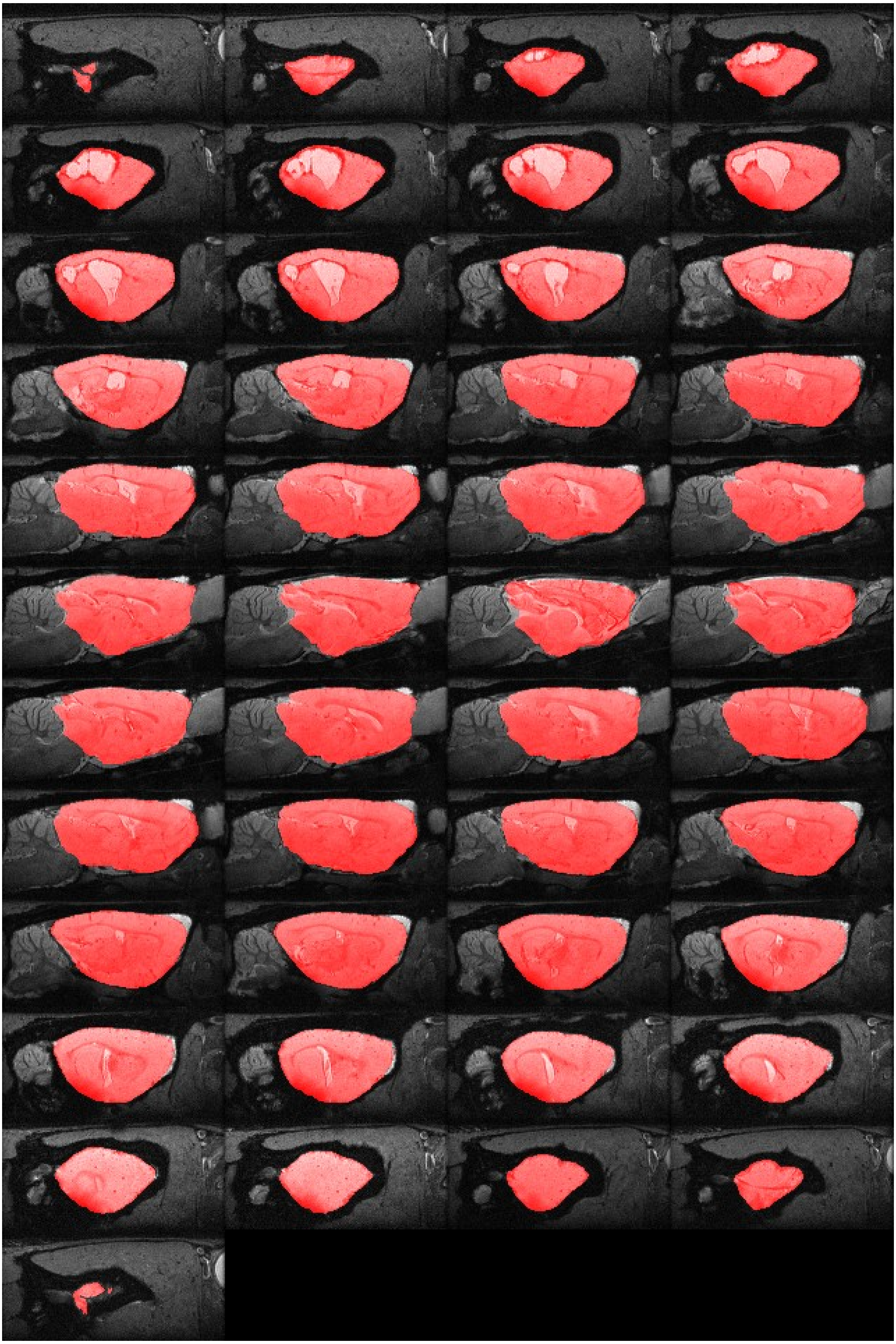

*Figure 2: Manual annotation of the brain mask in an EpiBioS4Rx rat, MRI scan acquired 150 days after TBI, displaying every annotated sagittal slice.*

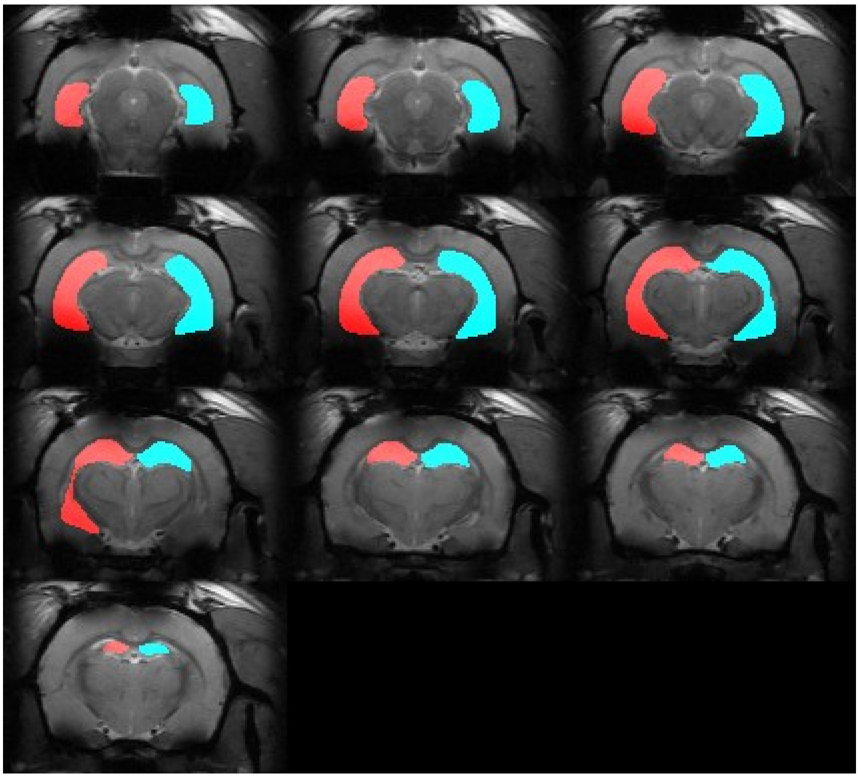

*Figure 3: Manual annotation of the hippocampus ipsilateral (red) and contralateral (blue) to the lesion, in an EPITARGET rat two days after TBI.*

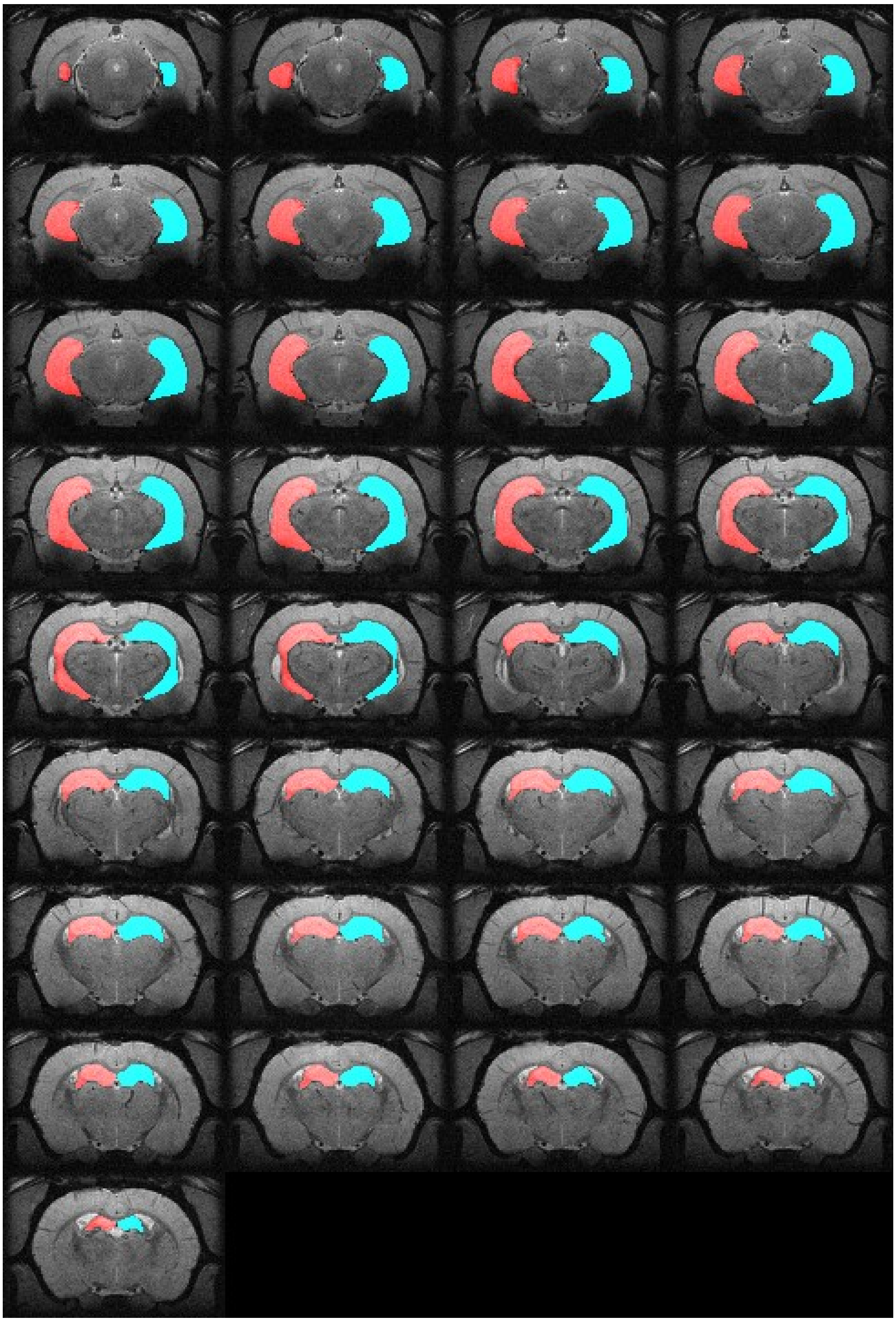

*Figure 4: Manual annotation of the hippocampus ipsilateral (red) and contralateral (blue) to the lesion, in an EpiBioS4Rx rat nine days after TBI.*

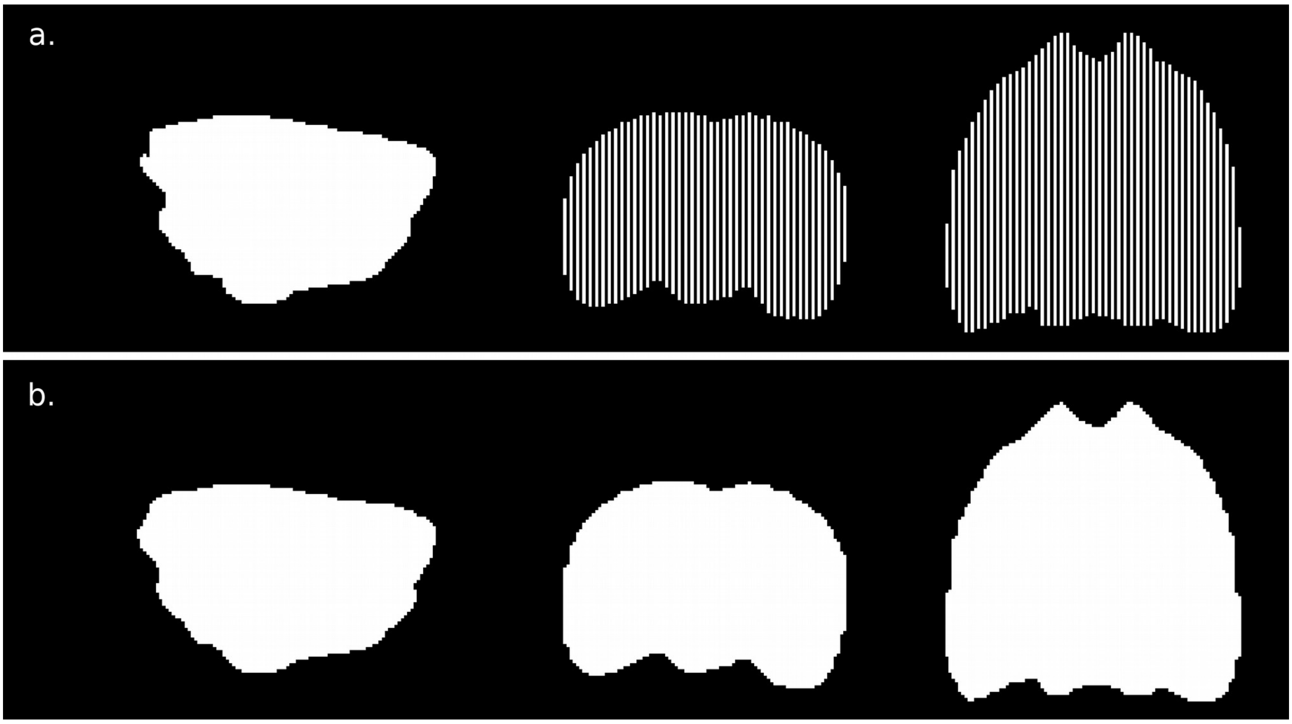


*Figure 5: Brain mask completion for the EpiBioS4Rx dataset. a. First, we manually labeled the brain mask in every second sagittal slice. b. Second, we applied a binary-closing operation to obtain a brain mask for the whole brian.*

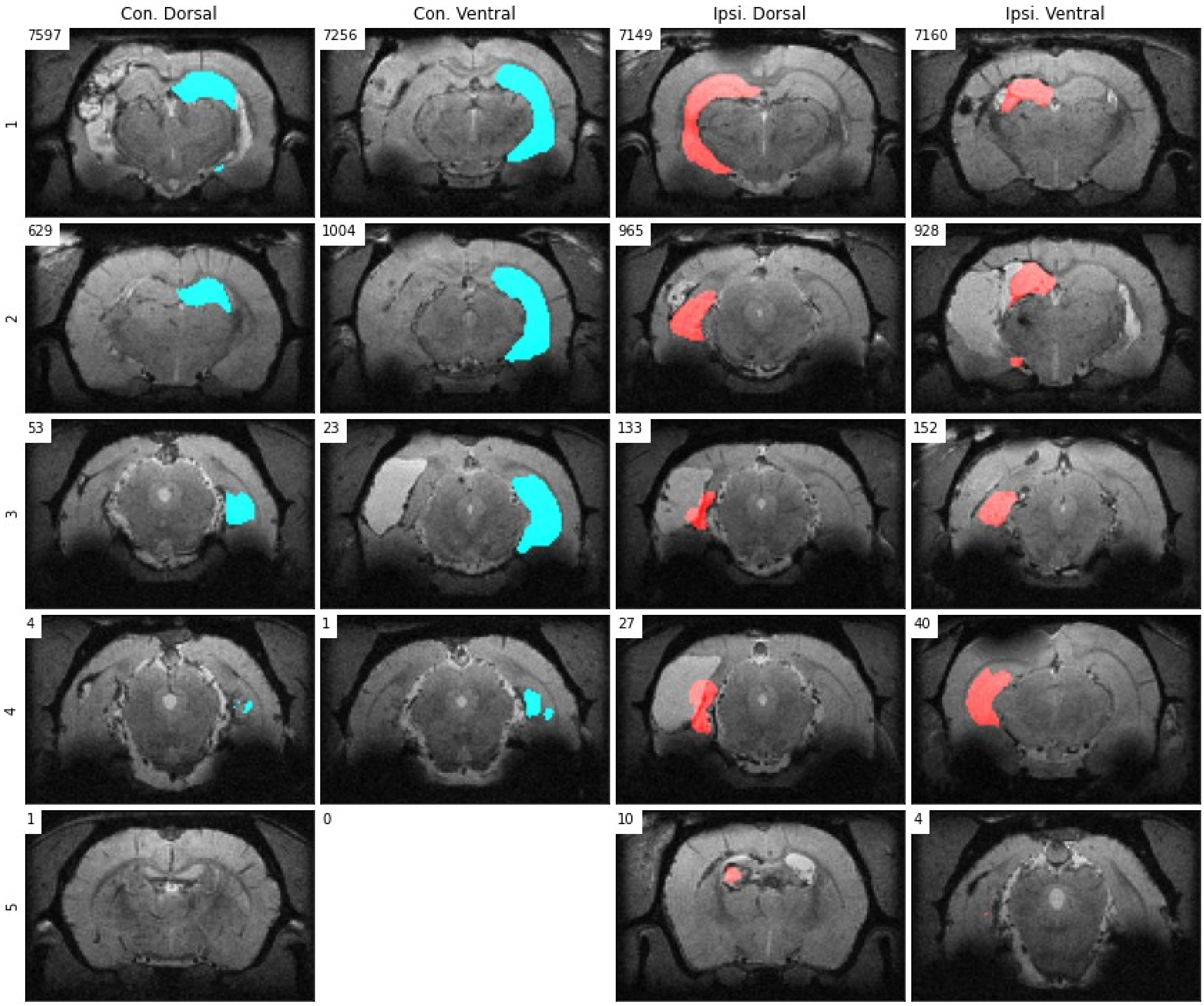


*Figure 6: Examples of MU-Net-R hippocampus segmentations and their visual evaluation scores, for the dorsal and ventral regions of the ipsilateral and the contralateral hippocampus, respectively outlined in red and blue. Numbers indicate the total number of slices annotated with each score in each region.*
